# Antibiotic uptake across gram-negative outer membranes: better predictions towards better antibiotics

**DOI:** 10.1101/667006

**Authors:** Ricardo J. Ferreira, Peter M. Kasson

**Affiliations:** Science for Life Laboratory, Department of Cell and Molecular Biology, Uppsala University, 75124 Uppsala, Sweden; Departments of Biomedical Engineering and Molecular Physiology and Biological Physics, University of Virginia, Charlottesville, VA 22908 USA

**Author notes:** . Address: Box 800886, Charlottesville VA 22908 USA.

## Abstract

Crossing the gram-negative bacterial membrane poses a major barrier to antibiotic development, as many small molecules that can biochemically inhibit key bacterial processes are rendered microbiologically ineffective by their poor cellular uptake. The outer membrane is the major permeability barrier for many drug-like molecules, and the chemical properties that enable efficient uptake into mammalian cells fail to predict bacterial uptake. We have developed a computational method for accurate prospective prediction of outer-membrane uptake of drug-like molecules, which we combine with a new medium-throughput experimental assay. Parallel molecular dynamics simulations are used to successfully and quantitatively predict experimental permeabilities. For most polar molecules we test, outer membrane permeability also correlates well with whole-cell uptake. The ability to accurately predict and measure outer-membrane uptake of a wide variety of small molecules will enable simpler determination of which molecular scaffolds and which derivatives are most promising prior to extensive chemical synthesis. It will also assist in formulating a more systematic understanding of the chemical determinants of outer-membrane permeability.

## INTRODUCTION

Antibiotic-resistant bacterial infections are an increasing threat to human health; if this trend is not reversed, it has been estimated that by 2050 drug-resistant infections will kill more people than cancer [1, 2]. In addition to better diagnostics and antimicrobial stewardship, development of new effective antibiotics is a critical part of the public health response. One challenge in developing antibiotics against gram-negative pathogens is small-molecule uptake: gram-negative membranes differ substantially from those of mammalian cells, and the heuristics that guide small-molecule selection for uptake in human cells are violated by most clinically approved antibiotics [3]. Effective uptake into gram-negative bacteria indeed often poses a barrier to preclinical development of otherwise promising antimicrobial leads. The gram-negative outer membrane provides a primary permeability barrier, and many antibiotics act in the periplasmic space between membranes, so outer-membrane uptake of small molecules is a good target for initial optimization of drug uptake in gram-negative bacteria.

Gram-negative bacteria have an outer membrane with a high proportion of lipopolysaccharide (LPS), and the majority of successful antibiotics and hydrophilic metabolites enter this membrane through nonspecific or moderately specific outer membrane porins [4,5]. Considerable effort has been expended developing experimental assays of small-molecule uptake either in aggregate or specifically through these porins [6–13]. Similarly, a number of statistical, and computational approaches have been developed to better understand the nature of small-molecule uptake through porins [14–20]. While these have provided substantial advances, they are not yet at a stage where they are routinely used in most antimicrobial development programs [21,22]. There is thus a need to further develop predictive and measurement approaches that can scale to help predict the uptake of new small-molecules in gram-negative bacteria.

We present here two approaches, one experimental and one computational, to improved prediction of small-molecule uptake across the gram-negative outer membrane. The experimental assay leverages prior work using liposome swelling as a measure of small-molecule uptake [7] but uses bacterial outer membrane vesicles (OMVs) to measure uptake across the bacterial outer membrane. The computational approach employs high-throughput molecular dynamics simulation to predict uptake in a quantitatively accurate, reproducible, and automated manner.

We adapt the inhomogeneous solubility-diffusion (ISD) model that has previously been used to estimate permeation across lipid membranes without pores [23] for use in measuring permeability through a membrane containing porins. Our approach is briefly summarized below; more details follow in the Methods. An initial permeability path is generated for transit from bulk water into the porin, through the narrower pore region, and then into the periplasmic space using a single long-time simulation where the molecule is slowly pulled through the pore (Fig. 1). Snapshots from this initial permeability path are then used to initialize umbrella-sampling simulations where the potential of mean force and local diffusion coefficient are calculated at intervals along the path. The inhomogeneous solubility-diffusion formalism is then used to calculate the permeability coefficient.

**Figure 1.**
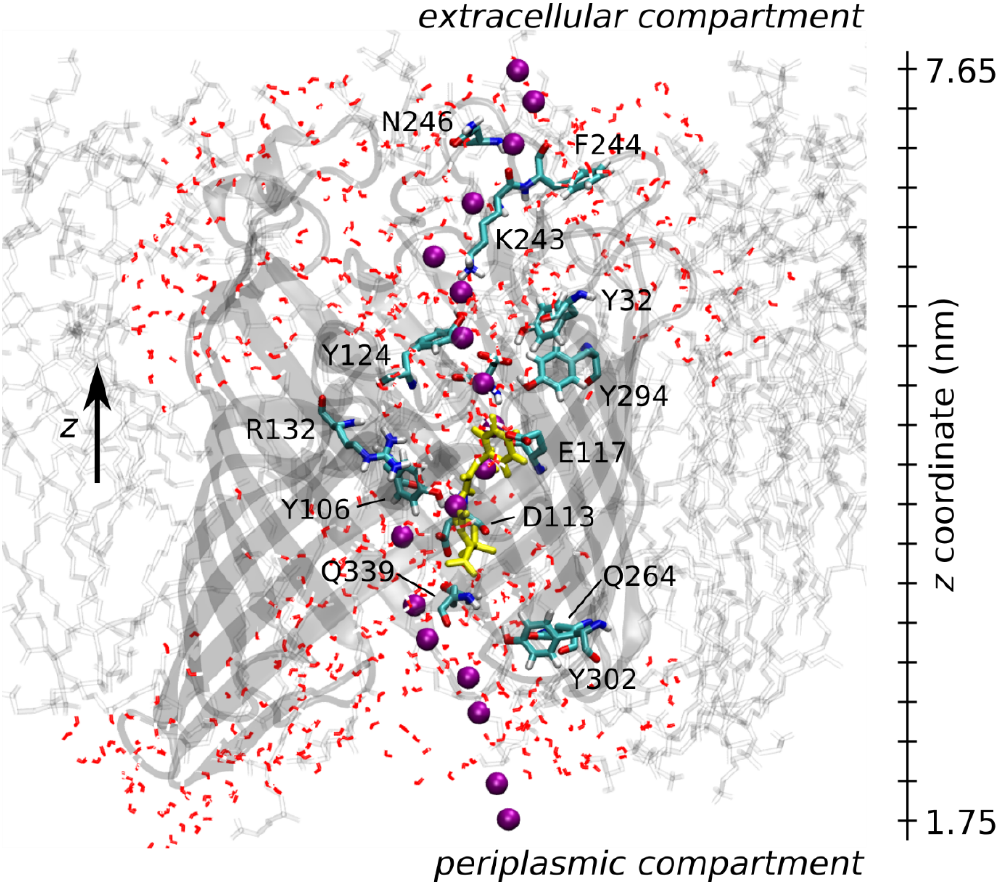
Simulation snapshot of the OmpF monomer, with some pore-aligning residues (along the generated path) and water molecules depicted in licorice. The starting position for umbrella windows 5-22 is represented by purple spheres (α-carbon at the β-lactam scaffold). An ampicillin molecule (yellow licorice) is also shown close to the constriction region.

In order to develop a simple, reproducible assay for measuring small-molecule uptake across the gramnegative outer membrane, we developed an assay that measures the osmotic swelling of small outer membrane vesicles derived from gram-negative bacteria (Fig. 2). Prior measurements of small-molecule permeability use one of several approaches: enzymatic detection of β-lactams entering the periplasmic space [8], calculating relative permeabilities into proteoliposomes with reconstituted porins via a liposome-swelling assay [7], measuring total compound accumulation in the cytoplasm via mass spectrometry [11] or electrophysiological measurements [24,25]. Indirect methods of measuring minimum inhibitory concentrations with and without alterations to porins or efflux pumps have also been used [26]. Of the direct measurement methods, liposome swelling assays measure permeability most directly; mass spectrometry provides an excellent detection method but is used for total cytoplasmic accumulation and thus takes into account both influx and efflux. However, liposome-swelling assays can be limited by the reconstitution procedure, including both membrane composition and asymmetry and porin insertional direction [27].

**Figure 2.**
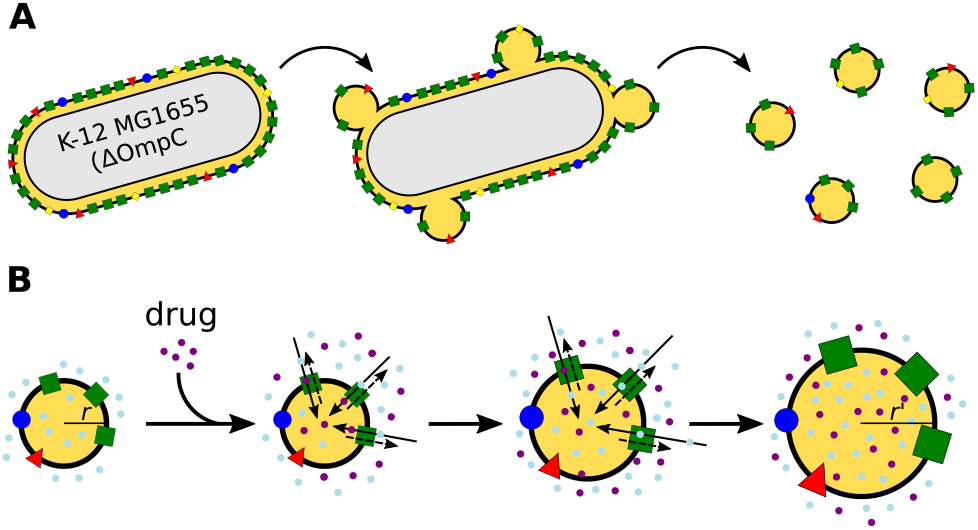
Preparation of outer membrane vesicles (OMVs). While from wild-type strains OMVs contain all major porins, OMVs from knockout strains are enriched in one major porin type (A). Isolated OMVs then undergo osmotic swelling in response to drug uptake.

In contrast, osmotic swelling of bacterial outer membrane vesicles provides a means to measure outer membrane permeability simply and directly in a bacterial membrane. Procedures for production and isolation of such OMVs have been well described [28,29], and they offer the advantage of a more physiological outer membrane (including LPS composition) than proteoliposomes [27]. Measuring osmotic swelling also permits the detection of a broader range of compounds than enzymatic detection approaches [8,30] and with less specialized apparatus than electrophysiological ones [24]. Drug permeability through specific outer membrane porins can further be measured using bacterial knockout strains deficient in the other porins, corrected for the relative abundance of the porins.

## RESULTS AND DISCUSSION

### Structural stability of OmpF simulated in E. coli outer membranes

The structure of the OmpF homotrimer remained stable over the “production” portion of the equilibration, with a mean RMSD value of 0.20 ± 0.01 nm (Fig. S1). Per-residue root mean-square fluctuation analysis showed the extracellular loops L6 (residues 240-250) and L8 (residues 320-330) as having the highest flexibility, as would be expected (Fig. S2). The position of the constriction loop L3 (residues 103-130) inside the water-filled pore was relatively stable throughout the whole simulation, and a slight decrease of the average volume of the pore chambers (6385 ± 1584 vs. 5897 ± 356 Å^3^, −8%) was also observed. The membrane structural parameters were relatively stable, with a slight increase in *A_L_* (218.5 ± 0.86 Å^2^ and 67.4 ± 0.28 Å^2^ for the upper and lower leaflets respectively) compared to the membrane system alone (200 ± 2.74 Å^2^ and 62.9 ± 0.86 Å^2^). No significant change was observed for *D_HH_* (35.2 ± 0.16 Å^2^ vs 35.2 ± 0.86 Å^2^ respectively). Both *A_L_* and *D_HH_* were found to be in agreement with experimental values or other published computational studies [31,32].

### Retrospective prediction of small-molecule uptake

We initially validated our simulation predictions using the inhomogeneous solubility-diffusion (ISD) model against previously published drug permeability data from liposome swelling assays and a published analytic scoring function that had been fit to the data previously [19]. Our model’s predictions show excellent agreement with the experimental data, with a slightly better Pearson correlation coefficient (*R*, 0.93 vs. 0.88) and lower root-mean square error (RMSE, 0.34 vs. 0.44) than the previous analytical model (Fig. 3). The prior scoring function is analytical and faster to compute but relies on previous knowledge on molecular properties and other parameters calculated using all-atom MD simulations (e.g. geometrical cross-section, internal electric field and electrostatic potential). Although computationally more involved, the method we report relies only on the umbrella sampling MD simulations without any additional parameters. Permeabilities predicted using the CHARMM force field were within the error range of those predicted using the GROMOS force field (Fig. S3).

**Figure 3.**
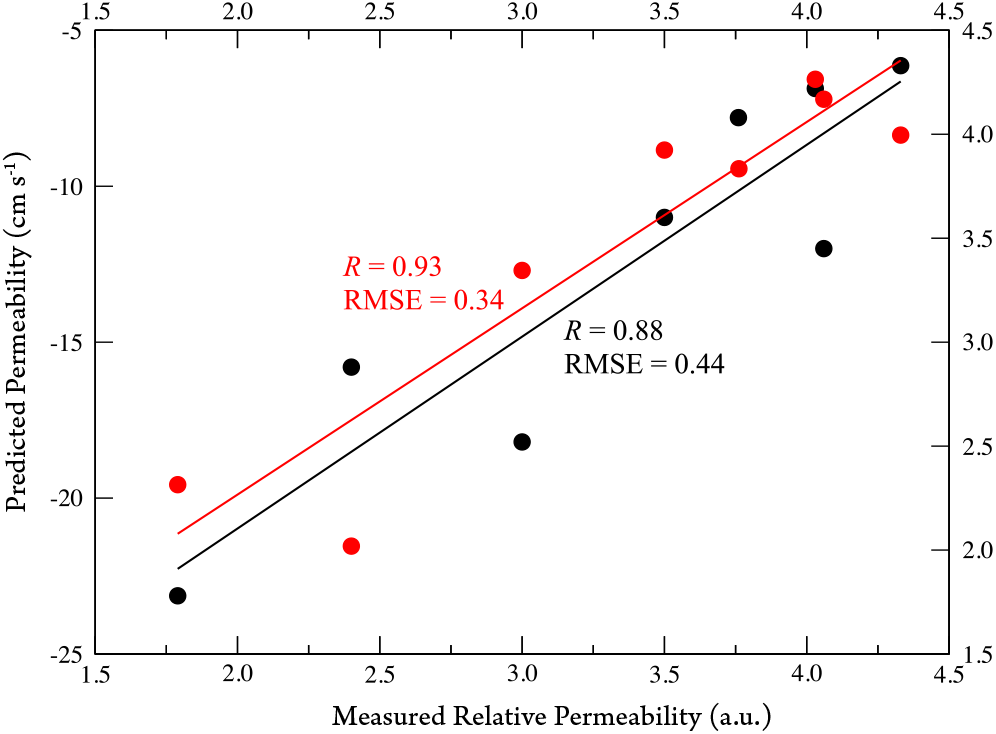
Retrospective validation of drug permeability. Predicted drug permeabilities are plotted against permeabilities previously measured via liposome swelling and compared against an analytical model fitted to those data and published accompanying them [19]. The analytical model is plotted in black and the new predictions using simulations and the inhomogeneous solubility-diffusion model in red. Permeabilities are normalized with respect to glycine as in previous work [19].

### Measurement of small-molecule uptake via outer membrane vesicles (OMV) swelling

After selecting 14 small molecules for prospective prediction and running the corresponding simulations, we measured outer-membrane permeability of these compounds through OmpF via swelling of OMVs derived from an *E. coli* MG1655 ΔOmpC strain. In the absence of added compound, the average radius of the isolated OMVs was 56.3 ± 10.3 nm and followed a log-normal distribution (Fig. S4). As in prior liposome-swelling assays, swelling rates for all compounds were normalized with respect to glycine (Fig. 4). Swelling was measured on crude OMVs without further purification, which improves scalability of the experimental assay but results in higher standard deviations (23 ± 4%). The characteristics of the harvested OMVs further depend on the incubation parameters but mostly on the growth phase in which they are harvested (ideally, at the late log growth phase). The standard deviations of measurements from a single OMV preparation are substantially lower.

**Figure 4.**
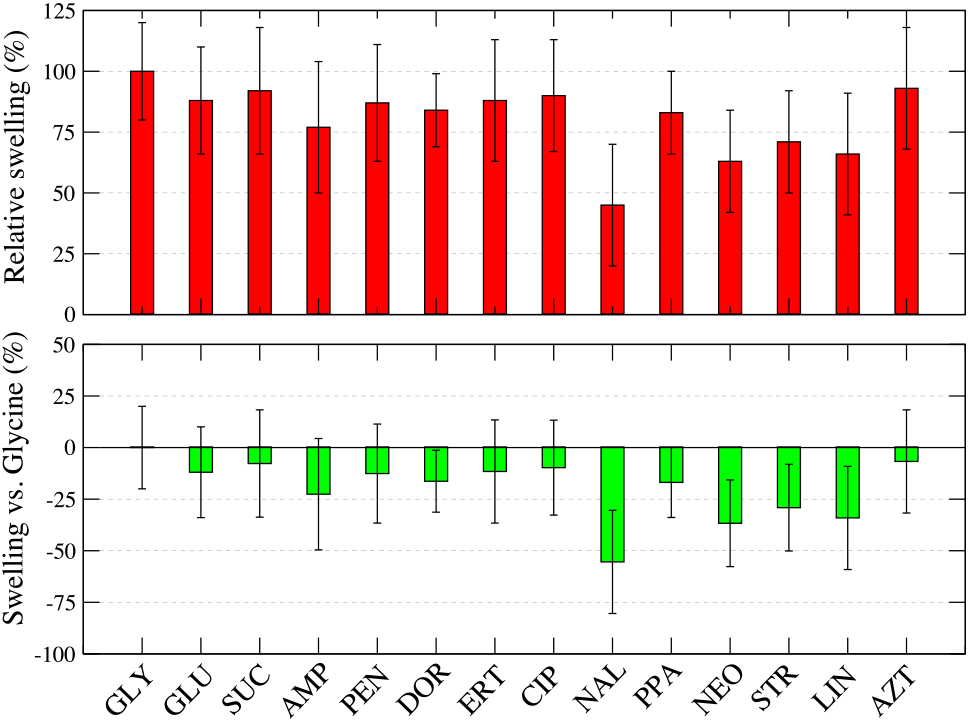
Relative permeabilities through OmpF-containing outer membranes. Permeation rates were determined from swelling experiments in ΔOmpC OMVs and measured via DLS. Data are plotted as mean ± standardization for the relative permeability of glycine and 13 other small molecules: glucose (GLU), sucrose (SUC), ampicillin (AMP), benzylpenicillin (PEN), doripenem (DOR), ertapenem (ERT), ciprofloxacin (CIP), nalidixic acid (NAL), pipemidic acid (PIP), neomycin (NEO), streptomycin (STR), linezolid (LIN) and aztreonam (AZT).

The resulting data reproduce at least the qualitative trends for previously reported permeabilities measured via other methods [6,7]. The four lowest-permeability compounds were nalidixic acid (45 ± 25%), linezolid (66 ± 25%) and the aminoglycosides neomycin (63 ± 21%) and streptomycin (71 ± 21%). Aztreonam (monobactam, 93 ± 25%), ciprofloxacin (quinolone, 90 ± 23%) and ertapenem (carbapenem, 88 ± 25%) were the antibiotics with higher relative diffusion rates. Glucose showed a slightly lower diffusion (88 ± 22%) than sucrose (92 ± 26%), which is in agreement with previous studies [9,33] but nonetheless still lower than those reported for the above antibiotics. Interestingly, benzylpenicillin displayed a higher relative diffusion rate (87 ± 27%) than ampicillin (77 ± 27%) [24], which may explain the high effectiveness of penicillin G.

### Correlation between predicted permeabilities and relative OMV swelling rates

Since prospective prediction provides the strongest test of a new method, we first simulated and then measured the outermembrane permeabilities of 14 small molecules. The predicted permeabilities showed a high Pearson correlation coefficient (*R* = 0.97) with measured OMV swelling rates (Fig. 5), following the well-established logarithmic relationship between permeability and swelling rate. For the quinolone series, nalidixic acid was predicted and measured to have a very low permeability (2.90E-11 cm s^−1^) while pipemidic acid and ciprofloxacin were predicted to have low (7.76E-06 cm s^−1^) and moderate (1.45E-04 cm s^−1^) permeabilities. These predictions agree well with the IC_50_ and MIC values reported in *E. coli* [34] and suggest that permeation through porins may be the limiting step on the quinolone mode of action in Gram-negative bacteria.

**Figure 5.**
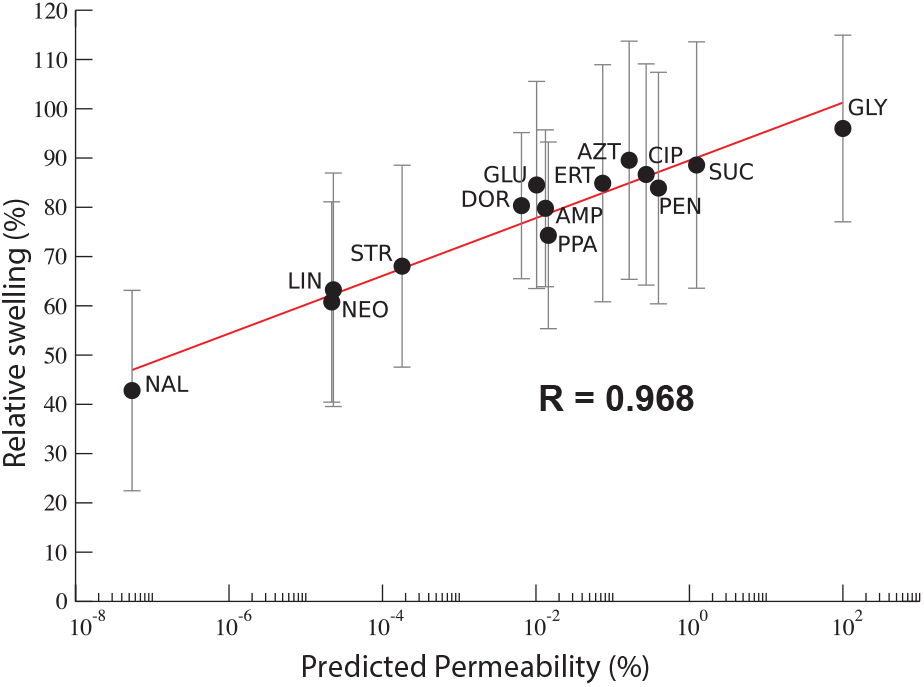
Prospective prediction and testing of small-molecule permeation through *E. coli* outer membranes containing OmpF. Plotted are semi-logarithmic correlations plots of experimental OMV swelling rates and predicted permeabilities of glycine (GLY), sucrose (SUC), benzylpenicillin (PEN), ciprofloxacin (CIP), aztreonam (AZT), ertapenem (ERT), pipemidic acid (PPA), ampicillin (AMP), glucose (GLU), doripenem (DOR), streptomycin (STR), linezolid (LIN), neomycin (NEO) and nalidixic acid (NAL).

Results for other compounds tested can also be well explained by their microbiological activities. Neomycin (1.16E-08 cm s^−1^), streptomycin (9.59E-08 cm s^−1^) and linezolid (1.22E-08 cm s^−1^) were predicted with very low permeabilities, but while aminoglycosides can penetrate the OM through a self-promoted pathway [35] linezolid is unable to penetrate the OM and does not have any substantial activity against most Gram-negative organisms [36]. Interestingly, the dianionic monobactam aztreonam was predicted and measured to have a moderate permeability through OmpF (8.72E-05 cm s^−1^). This is not in agreement with the diffusion rates previously published from liposome-swelling assays but provides a better correlation with reported IC_50_ and MIC values in *E. coli* strains [37,38]. The permeabilities of the carbapenems doripenem (3.44E-06 cm s^−1^) and ertapenem (3.94E-05 cm s^−1^) also well explain the published microbiological activities of these compounds [39,40].

Several compounds had surprising results: while glycine was, as expected, predicted to have the greatest permeability through OmpF (5.32E-02 cm s^−1^), glucose was measured to have lower permeabilities through OmpF (5.58E-06 cm s^−1^) than sucrose (6.69E-04 cm s^−1^) or lactose (3.01E-03 cm s^−1^), which is not the case in whole cell assays [9]. Here, our predictions are consistent with our OMV swelling measurements: the difference between glucose and sucrose is greater in our simulations though within experimental error. However, this may be a question of uptake mechanism: glucose can cross the outer membrane via the maltose outer membrane LamB porin while sucrose and lactose do not. Since the *E. coli* strain used for experimental measurements was grown in the absence of maltose, expression of LamB should be low. Also, within the penicillin series, while ampicillin (8.75E-06 cm s^−1^) is within the same range as experimentally measurements in intact cells [41], benzylpenicillin was shown to have higher permeabilities (2.11E-04 cm s^−1^) but nonetheless still in agreement with a previous study [24] that suggested a different interaction mode with OmpF.

We further analyzed the calculated potential of mean force (PMF) profiles to better understand why glucose was predicted to have lower permeability than sucrose. The free-energy barrier for glucose transit through OmpF is higher than for sucrose (100 vs. 60 kJ mol^−1^, respectively) (Fig. S5A), with a local minimum in the preorientation chamber (*z* ~ 5.0 nm) and the energy barrier spanning the entire constriction zone up to the porin-lipid interface at the periplasmic side (*z* ~ 3.0 nm). In contrast, sucrose did not display any substantial free-energy minima, and the only meaningful barrier to permeation is the constriction zone (Fig. S5B).

One of the first studies using intact cells obtained higher permeation rates for hexoses than for disaccharides [9], although it was not determined which porins were responsible. Our results therefore suggest that although OmpF may not be the preferred entry route for glucose (but instead the maltose LamB transporter), it may be an important permeation route for other sugars as sucrose or lactose.

We performed a similar analysis to why ampicillin was predicted to have lower permeability than penicillin. Despite similar free-energy maxima (~80 kJ mol^−1^), the PMF for ampicillin shows permeation to be mostly unfavorable, with an overall *ΔG* of +40 kJ mol^−1^ (Fig. S5C), while for benzylpenicillin the overall *ΔG* was −20 kJ mol^−1^ (Fig. S5D). Geometrically, while ampicillin dipole-dipole interactions between the positively charged amino group with the residues L115, P116 and E117 of the L3 loop and between the negatively charged portions of ampicillin with the positively charged residues R42, R82 and R132 were observed, for benzylpenicillin only the latter were identified. Although the presence of charged amino groups is expected to promote accumulation within the periplasm [17], our results demonstrate that non-hindered charged groups such as primary or secondary amines may also impair drug permeation through the OmpF porin.

### Comparison to whole-cell permeability measurements

We also evaluated computational predictions of small-molecule permeability against whole-cell measurements for 19 compounds reported by Nikaido *et al.* [8,9,42] and Matsumura *et al*. [43]. Our calculated OmpF permeabilities were within whole-cell experimental error [43,44] for 11 out of 19 compounds (58%), with additional 3 compounds predicted within the experimental range but less accurately (16%). For the remaining compounds– SCE-20, cephapirin, cephalothin, piperacillin and benzylpenicillin (26%)– the predicted permeabilities through OmpF did not agree with whole-cell uptake measurements (Table S1), perhaps due to finite-box-size electrostatic effects [10,45].

Given the differences in assay system between whole-cell permeability and OmpF transit, this is remarkably good predictive power. There are two experimental reasons for the observed disagreements, even neglecting any possible discrepancies between predicted OmpF permeability and measured OmpF permeability. First, the experimental measurements use several different *E. coli* strains, many of which were engineered for high β-lactamase expression (either of a native or an exogenous enzyme), in order to facilitate enzymatic detection. In the dataset we used, the following strains were used: *E. coli* K-12, *E. coli* MC4100/pSW313 expressing a group 3 β-lactamase, *acrAB* deletion mutant of *E. coli* LA51, *E. coli* overexpressing 10-fold *ampC* β-lactamase and *E. coli* LGC10 carbapenem-hydrolyzing βlactamase. This can of course affect permeability values.

Second, whole-cell compound accumulation involves several other drug transport mechanisms, including the presence of other porins, specific transporters and efflux pumps activity [4]. Several of the compounds measured have different substrate profiles for the AcrB efflux pump [46]. While cephaloridine is a ‘border-line’ substrate with high *V*_max_ (1.82 nmol mg^−1^ s^−1^) with an high *K*_0.5_ (288 μM), cephalothin was found to be a much better substrate, with *V*_max_, *K*_0.5_ and *k*_cat_ of 1.0 nmol mg^−1^ s^−1^, 91.2 μM and ~500, respectively. Although the same study reported cefazolin not to be an efflux pump substrate, both cefazolin and cephalothin were found to have the lowest *K*_M_ for R_471a_ β-lactamase in the original study [8], which can also explain the previously reported lower permeabilities.

## CONCLUSIONS

The development of antibiotics is currently impaired by the low success rate in translating new biochemical hits into approved drugs. Methods to improve small-molecule uptake in gram-negative bacteria would help alleviate one substantial bottleneck in this process. Here, we report a simulation-based method to estimate drug permeability coefficients through the OmpF porin, the most important entry route providing access to the periplasmic space in bacteria. Results from this method show excellent agreement with experimentally measured permeability coefficients when performed in the same strain. We also report the development of an outer-membrane-vesicle swelling assay to specifically and scalably measure small-molecule permeation across the bacterial outer membrane. This could be used to assess the impact that chemical modifications would have on the permeability of a given hit or lead compound prior to the synthesis of such modifications.

The computational approach described here has several practical advantages. It uses simple simulation methods easily available in multiple MD packages. Our calculations took 1.1 *μ*s of simulation time and <1 week of wall-clock time per molecule and could be run in parallel on all molecules simultaneously. We also benchmarked our calculations using Google Compute Engine in June 2018; the approximate cost of predicting a small molecule permeability was $1312 without extra discounts. Our approach permits further automation (without modifying the underlying method) for large-scale deployment using high-level simulation interfaces such as *gmxapi* [47] on either clusters or cloud-computing platforms. The goal of this would be twofold: large-scale deployment and elimination of any manual tuning for complete reproducibility.

Currently, apart from appropriate ligand parameterization, the primary need for expert assessment is in setting the pulling rate. Faster pulling is more computationally efficient, but the pull rate must allow some degree of ligand equilibration within the porin, and fast pulling can induce conformational changes in the constriction region.

We have presented a new experimental assay using outer membrane vesicles (OMVs) to determine antibiotic permeation through porins. OMVs have the advantage of retaining the composition of the outer membrane, and in particular the permeability and interfacial differences conferred by lipopolysaccharide. This assay can be used in conjunction with clinical isolates, model strains, or knockout strains that predominantly express a single porin species. OMVs are naturally shed by bacteria and can be produced in a matter of hours and are thus faster to prepare than proteoliposomes. The dynamic light scattering assay can further be performed in 96-well format.

Bacterial strain and growth conditions can substantially impact permeability of the outer membrane and cytoplasmic accumulation even more so. In whole-cell uptake studies, Matsumura and co-workers [43] obtained permeabilities for imipenem and meropenem that were 4- to 5-fold lower than those obtained in a previous study [44]. While Matsumura’s paper suggests that this difference may be due to the very weak barrier of the outer membrane in *E. coli* LCG10 used by the other study, this difference may also be due to the different *K_M_* values of the β-lactamases used for substrate detection (*K_M_* of 962 and 281 μM for imipenem and meropenem, respectively versus 42.7 and 4.75 μM). Furthermore, a recent study on *Enterobacter cloacae* also revealed that AmpC overproduction may be associated with *ompF* gene downregulation, a mechanism that could also contribute to the lower permeabilities reported for ampicillin and benzylpenicillin [48]. Since these results are intimately dependent on the used *E. coli* strain, the type and/or expression of β-lactamases, bacterial strains should be selected carefully for optimal relevance. Our computational approach and OMV swelling assays of course do not avoid this problem, but they do permit measurements on more precisely defined membranes that can then be calibrated against the desired conditions.

## METHODS

### Selection of experimental data

Compounds previously measured via whole-cell uptake assays were compared retrospectively. Permeability coefficients (in cm s^−1^) were determined in intact cells [8,41,43] and included twelve cephalosporins [8,43,49], three penicillins and two carbapenems [41,43]. Lactose [9] permeability in *E. coli* B (CM6 strain) was also included in this study to compare the permeation rates of neutral molecules through OmpF with the ones obtained from antibiotics.

A second set of compounds was selected for prospective prediction of permeability followed by experimental measurement. Several classes of small molecules are represented, among which aminoacids (glycine), sugars commonly used to standardize uptake measurements (glucose, sucrose), penicillins (ampicillin, benzylpenicillin), carbapenems (doripenem, ertapenem), quinolones (nalidixic acid, pipemidic acid, ciprofloxacin), aminoglycosides (neomycin, streptomycin), a monobactam (aztreonam) and an oxazolidinone (linezolid).

### System construction for molecular dynamics simulations

An outer-membrane patch (OM) containing 120 lipopolysaccharides (LPS) in the upper leaflet and 381 lipids in the lower leaflet [276 1-palmitoyl-2-oleoyl-*sn*-glycero-3-ethanolamine (POPE), 68 1-palmitoyl-2-oleoyl-*sn*-glycero-3-phosphoglycerol (33 *D*-*isomer* and 35 *L*-isomer, respectively) and 37 cardiolipin (1,3-bis(*sn*-3’-phosphatidyl)-*sn*-glycerol, POPO)] was constructed in GROMACS v2016.4, following previous studies by Piggot *et al.* [50,51]. As the available parameterization for the inner core of LPS only included two keto-deoxyoctulosonate (KDO) residues, the parameterization for three additional heptoses (L-glycero-D-*manno*heptulose), two of which phosphorylated, was obtained in PRODRG server [52] and the acyl chains in lipid A readjusted to match the lipid parameters by Poger *et al.* [53,54]. Following the approach taken in previous reports, Ca^2+^ and Mg^2+^ were added to the membrane to neutralize the repulsion between charged LPS molecules while Na^+^ was used for neutralizing the remaining charges. A similar membrane patch (175 LPS, 350 POPE, 100 POPG and 50 POPO lipids) was obtained for the CHARMM36m force field using CHARMM-GUI [55–57]. Each membrane patch was then equilibrated in GROMACS for at least 100 ns of simulation time, after which the total area per lipid (A_*L*_) and thickness (*D_HH_)* were checked for consistency with experimental data [31,58].

The initial structure of *E. coli* outer membrane porin F (OmpF) was taken from crystallographic data (PDB ID: 3K19) [59]. Protonation states were assigned using PROPKA 3.0 [60,61], the structure was converted to the GROMOS 54a7 force field and inserted in the previously equilibrated OM patch using *g_membed* [62,63]. perpendicular to the membrane *xy* plane and orienting the longer extracellular coils towards the LPS outer leaflet. The system was also prepared using the CHARMM36m force field as noted below. The whole system (protein and membrane patch) was then further equilibrated via energy minimization, a 100 ps *NVT* equilibration run at 310 K (above the lipid gel-fluid transition temperature) [64], and a 25 ns *NPT* run to allow adjustment between the membrane patch and OmpF, keeping all non-hydrogen atoms of the homotrimeric protein spatially restrained. After this, the position restraints were removed in a step-wise fashion over two 10-ns simulations: first all but the backbone and second all but Cα, C, and N atoms were released. A 100-ns unrestrained simulation was then performed to assess the structural stability of system, with the last 50 ns used for analysis of these properties. Membrane properties as A_*L*_ and thickness *D_HH_* were re-assessed, and the average volume of the water-filled pores was calculated using CavitySearch (http://eds.bmc.uu.se/eds/cavitysearch.php) or estimated from the number of water molecules found inside and in-between the phosphate headgroups of the upper and lower leaflets.

### Small molecule parameterization

All molecules except glucose were parameterized according to the GROMOS96 force field in PRODRG online server [52] and manually curated, with partial charges assigned using the AM1-BCC charge model [65,66] in Antechamber 1.27 [67], as previously suggested [68]. Glucose was parameterized according to the 56A6_CARBO_ force field [69]. For usage in CHARMM systems, molecule topologies were obtained using the CGenFF server (https://cgenff.umaryland.edu) [70,71] and converted to GROMACS format using the cgenff_charmm2gmx.py script. Parameter sets are available from the accompanying Dryad package.

### Estimation of initial permeability paths using pulling simulations

A single molecule was placed above the first OmpF monomer using the end state of the final 100-ns run above. The molecule was placed in bulk water, at roughly ~1.2 nm from the protein, and any overlapping waters were removed. This molecule was then pulled to the opposite side of the OM patch along the *z*-axis (perpendicular to the membrane plane and parallel to the water-filled pore) over 100 ns, using a spring constant of 1000 kJ mol^−1^ nm^−2^ and a pull rate of 0.085 ± 0.005 nm ns^−1^. This pull rate was chosen to allow enough time for the protein to adjust to the presence of the molecule passing through its internal pore. Simulation parameter files are available in the accompanying Dryad package.

This trajectory was used to obtain an initial permeation path, and starting points for umbrella sampling were chosen at even *z*-intervals along this path. For all molecules, 26 umbrella windows with a mean width of 0.35 ± 0.02 nm were used to sample the *z* coordinate path with at least 40 ns of MD was performed per window (>1 *μ*s total sampling per molecule). The convergence of the PMF was assessed by generating PMFs at different umbrella-sampling intervals; if convergence was inadequate at 40 ns, the umbrella-sampling runs were extended to 100 ns.

### Prediction of permeabilities through OmpF

The inhomogeneous solubility-diffusion (ISD) model was used to calculate drug permeabilities via the following relation:

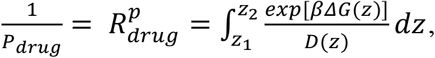

in which *β* = 1/*k*_*B*_*T*, Δ*G(z)* is the potential of mean force (PMF), *D(z)* is the local diffusivity coefficient and *z* is the relative position of the molecule along the pore axis. Both Δ*G(z)* and *D(z)* can be estimated from MD simulations if all *z* positions of the molecule are adequately sampled.

The local diffusivity coefficient *D(z)* was calculated from each umbrella window according to the approach derived by Hummer (following the work of Woolf and Roux) [72]:

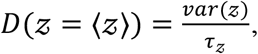

where ⟨*z*⟩ is the average of the coordinate path *z* in the biased run, *var*(*z*) = ⟨*z*^2^⟩ − ⟨*z*⟩^2^ is its variance and *τ*_*z*_ the autocorrelation time. We employ an approach developed by Zhu and Hummer [73] to estimate *τ*_*z*_ from the variance of the mean value *z* (estimated using the block averaging method by Flyvbjerg and Petersen) [74] and the interval of the time series data points (*Δt*):

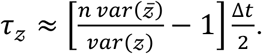

In estimating ⟨*z*⟩, we assume that the autocorrelation function *C(t)* vanishes at a time scale much shorter than the overall trajectory length *n*Δ*t*, and therefore the variance of *z̄* will also depend on *τ*.

The local resistance to permeation, and the permeability as its inverse, can thus be expressed using the above diffusivity relationship and the potential of mean force for each bin of the umbrella sampling. Error estimation was performed by bootstrap on the umbrella sampling windows sampled at intervals equal to the calculated autocorrelation times in each umbrella window. Full calculations for selected small molecules were also performed using the CHARMM36m force field.

### Isolation of outer membrane vesicles

Crude outer membrane vesicles (OMVs) were obtained as previously described [28]. 5 μL of an OmpC knockout strain (*E. coli* MG1655, ΔOmpC, kind gift of Linus Sandegren) was inoculated in 5 mL LB broth and incubated overnight at 37°C/200 rpm. This culture was then seeded into 250 mL of fresh LB broth and incubated for 4 h at 37 °C/100 rpm or OD_600_ between 1.3 and 1.4. This culture was centrifuged for 20 min at 10.000 ×*g* and 4 °C. The supernatant was then sterile-filtered through a 0.22 μm filter followed by ultra-centrifugation for 4 h at 150.000 ×*g*, 4 °C. The supernatant was immediately decanted, and the pellet was resuspended in 1 mL MES buffer pH 6.0, for DLS measurements.

### Dynamic light scattering (DLS) analysis

Each compound was dissolved in 100 μL of MES buffer pH 6.0 to a final concentration of 9 mM (charged molecules) or 18 mM (uncharged) [6,7]. In order to measure vesicle swelling, 6 μL of the OMV suspension was added to 6 μL MES buffer containing 2 μL MES (control) or 2 μL of the desired antibiotic, to a total volume of 14 μL. The autocorrelation function was measured over 60 seconds using a 660 nm laser on an AvidNano W130i instrument (Avid Nano, High Wycombe, UK). This function is the cross-correlation of the scattered intensity signal with itself, carrying information about the size of the particles because of the Brownian motion. If monodisperse, the autocorrelation function is a simple exponential decay with an exponent proportional to the hydrodynamic radius, but in the case of polydisperse samples the autocorrelation function becomes the sum of exponential decays, each corresponding to a subspecies of the distribution. All measurements were performed using a quartz cuvette and processed with iSize 2.1 (Avid Nano, High Wycombe, UK). At least 5 independent measurements were performed for each molecule.

## Supporting information

Supplementary Methods and Results

## ACKNOWLEDGMENTS

The authors thank L. Sandegren for providing bacterial strains, A. Villamil Giraldo and P. Purhonen for assistance with electron cryo-microscopy, and R. Nakamoto for many helpful discussions.

