## Supplementary Methods and Results for "Antibiotic uptake across gram-negative outer membranes: better predictions towards better antibiotics"

**Automated prediction of antibiotic uptake across gram-negative outer membranes**

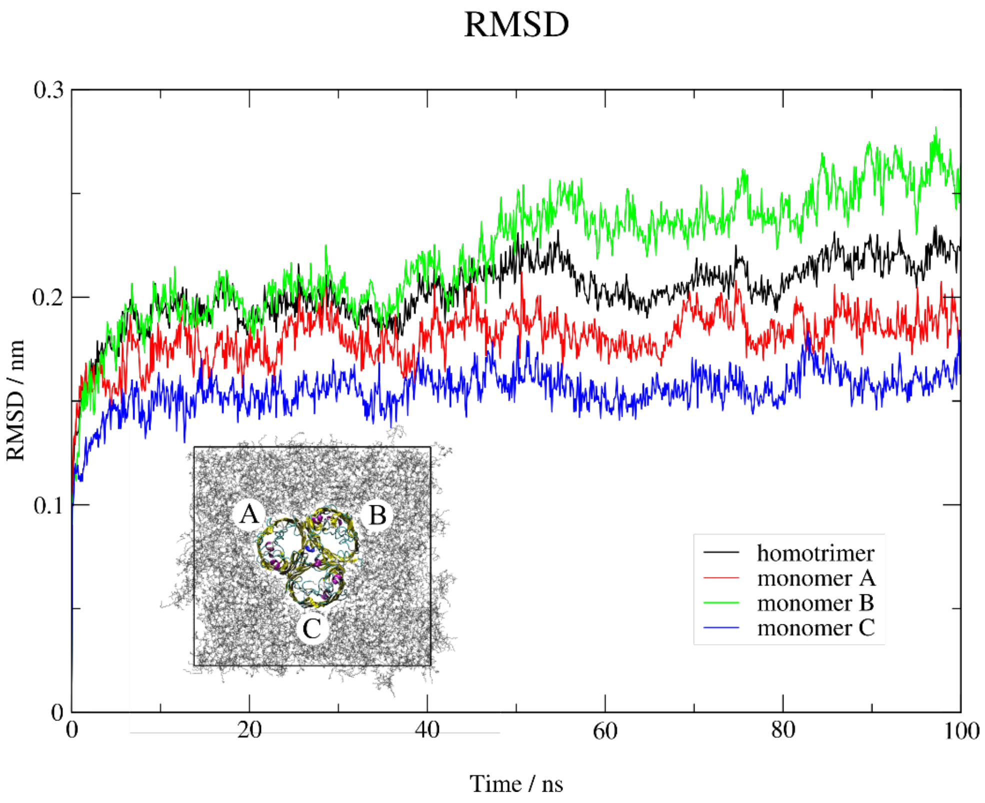


**Figure S1.** Root-mean square deviation (RMSD) for OmpF homotrimer and individual OmpF monomers during a 100-ns simulation.

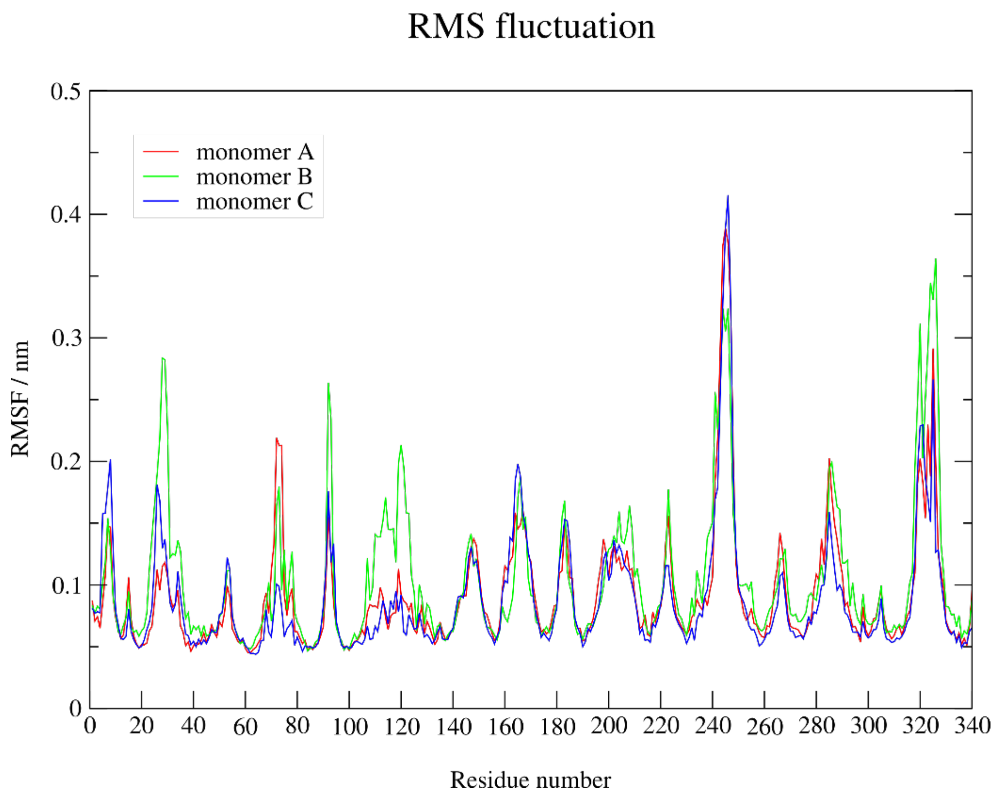


**Figure S2.** Root-mean square fluctuation (RMSF) of individual OmpF monomers during a 100-ns simulation.

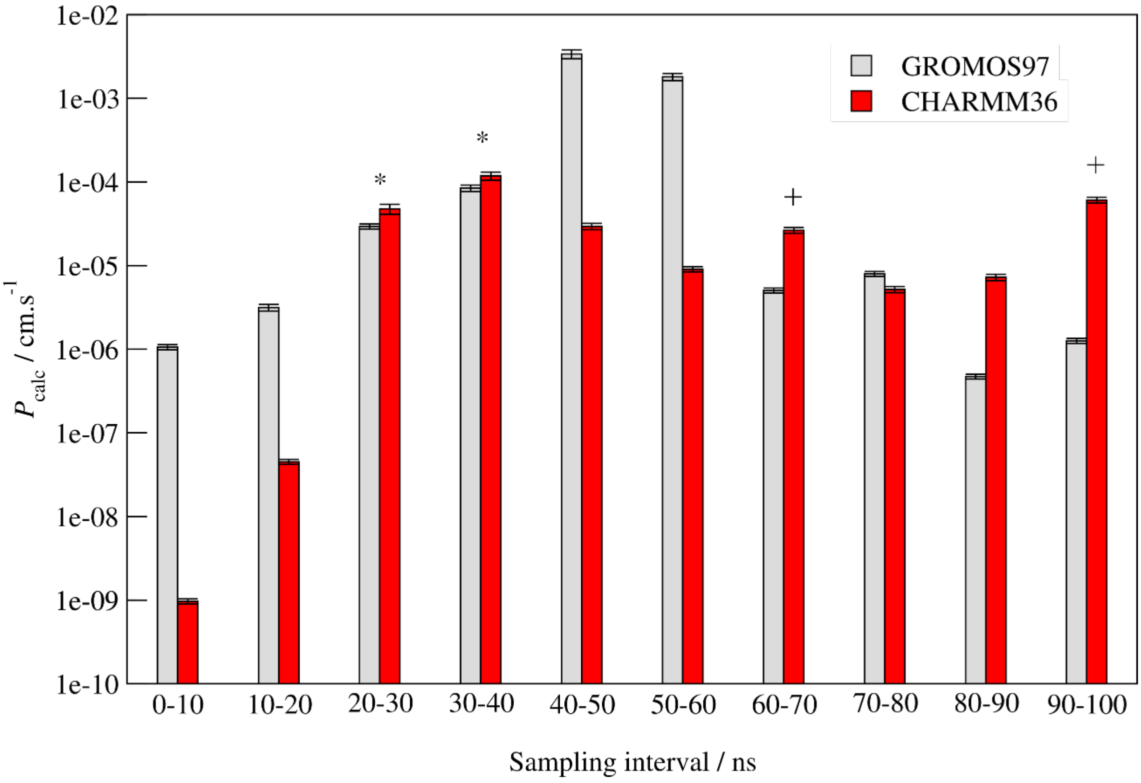


**Figure S3.** Extended 100-ns umbrella-sampling simulations for cephacetrile. Overall calculated permeabilities are 8.44E-05 ± 7.31E-06 $cm.s^{-1}$ with GROMOS97 and 11.8E-05 ± 1.33E-05 $cm.s^{-1}$) with CHARMM36. Annotations are as follows: * denotes results within experimental error for both force fields, while + denotes results within experimental error for CHARMM36 only.

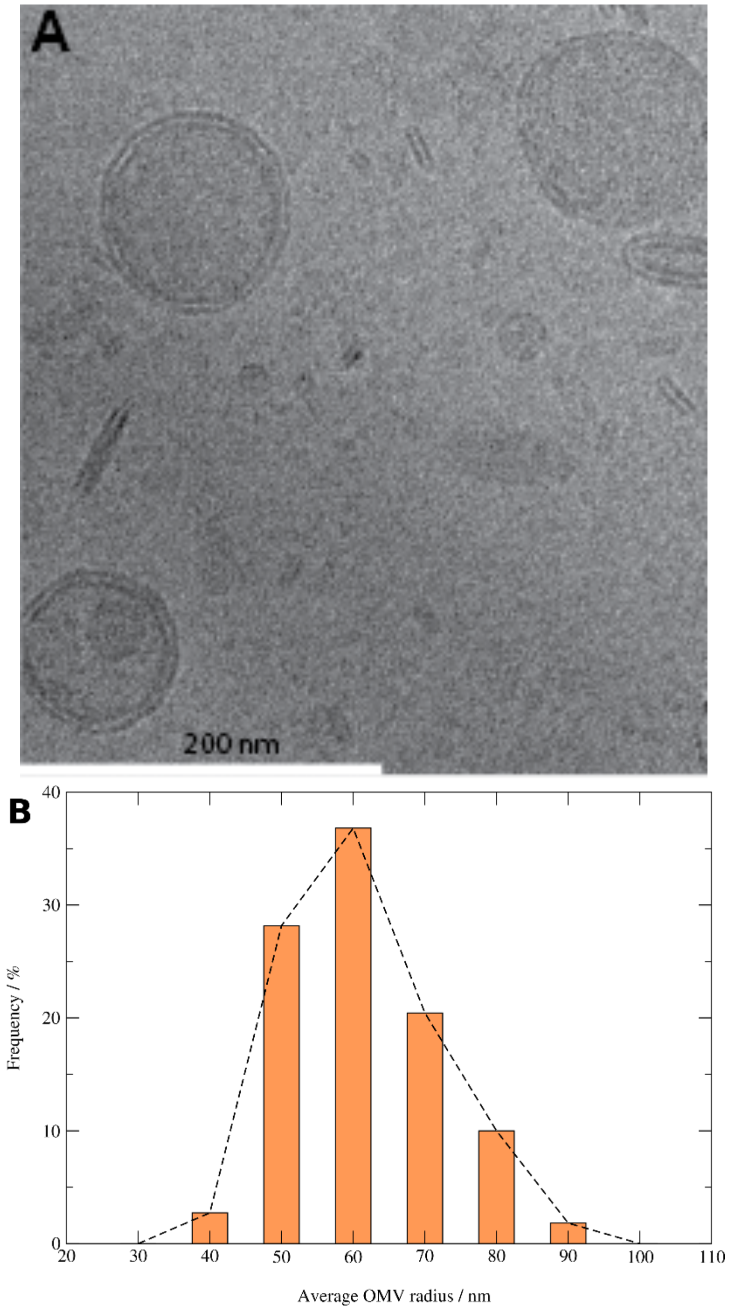


**Figure S4.** A) electron micrograph of outer membrane vesicle preparation and, B) size distribution for isolated OMVs. For cyo-electron microscopy, OMV samples were frozen on continuous carbon grids using a Vitrobot (FEI, Eindhoven, The Netherlands). 3 uL sample was pipetted to a glow-discharged, carbon-coated copper grid. The grid was blotted for 5s at 22 °C, ~90% humidity. Data were collected at 200 kV with a JEM-2100f microscope (JEOL Ltd., Tokyo, Japan) equipped with a TVIPS TemCam-F415 4k x 4k CCD camera. Micrographs were imaged at 50,000x magnification.

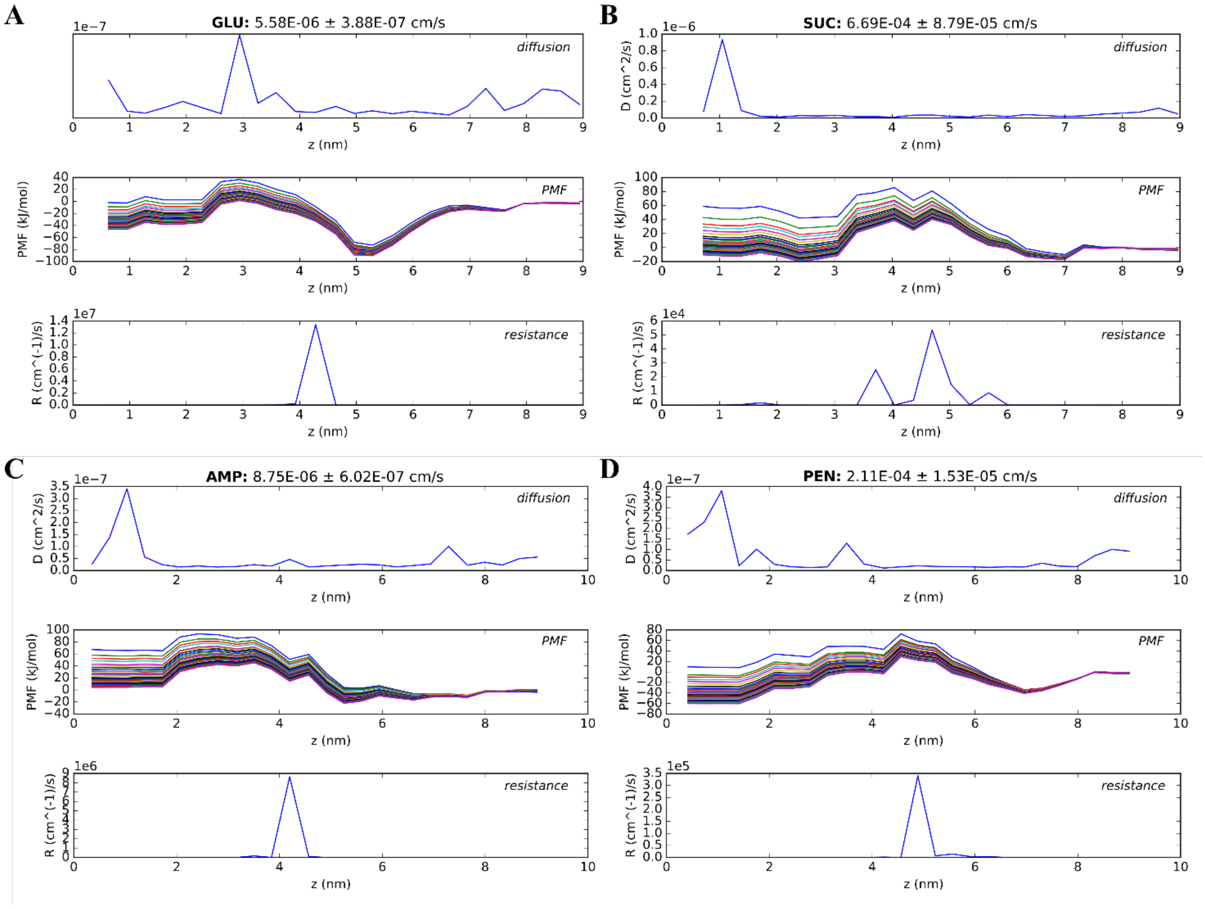


**Figure S5.** Analysis of calculated permeability profiles**.** Plotted are diffusion, potentials of mean force (different lines indicate bootstrap-resampling runs), and resistance to permeation profiles from permeability calculations for (A) glucose, (B) sucrose, (C) ampicillin and (D) benzylpenicillin.

**Table S1.** Predicted and previously measured whole-cell permeability values.

| compound | charge | MW | $P_{EXP}$ (x $10^{-5}$ cm.s$^{-1}$) | $P_{CALC}$ (x $10^{-5}$ cm.s$^{-1}$) |
|---|---|---|---|---|
| cephaloridine | + – | 415 | 53.7 | 18.3 ± 1.2 |
| imipenem | + – | 299 | 17.5 ± 2.0 * | 7.5 ± 0.5 |
| meropenem | + – | 383 | 3.0 ± 0.3 * | 1.2 ± 0.1 |
| ampicillin | + – | 349 | 0.28 ¥ | 0.77 ± 0.05 |
| cefpirome | + – | 514 | 0.7 ± 0.1 * | 0.2 ± 0.02 |
| cephaloglycin | + – | 406 | 9.8 | 1.9 ± 0.1 |
| | – | 405 | 4.7 | 0.45 ± 0.01 |
| cephapirin | + – | 421 | 19.4 | 0.30 ± 0.02 |
| | – | 420 | 3.2 | 1.13E-5 ± 1.0E-06 |
| cefazolin | – | 453 | 15.8 | 47.4 ± 5.8 |
| cephacetrile | – | 338 | 7.5 | 8.4 ± 0.7 |
| cephalothin | – | 395 | 2.90 | 21.9 ± 2.1 |
| cefamandole | – | 461 | 1.0 | 3.7 ± 0.2 |
| piperacillin | – | 516 | 2.7 ± 0.5 * | 1.7E-5 ± 1.1E-6 |
| cephaloram | – | 389 | 0.8 | 1.7 ± 0.1 |
| BTC | – | 445 | 0.2 | 0.09 ± 0.007 |
| penicillin G | – | 333 | 0.07 ¥ | 21.0 ± 1.5 |
| cefsulodin | + – – | 531 | 1.8 | 2.3 ± 0.2 |
| ceftazidime | + – – | 546 | 0.3 ± 0.1 * | 0.01 ± 0.001 |
| SCE-20 (Takeda) | – – | | 0.29 | 1.5E-04 ± 1E-05 |
| lactose | 0 | 342 | > 10 # | 301 ± 37.8 |

* evaluated in *E. coli* MC4100/pHS313 expressing group 3 β-lactamase; # evaluated in *E. coli* CM6; ¥, evaluated in *E. coli ΔacrAB::kan* LA51A. All remaining molecules were evaluated in *E. coli* K-12 $R_{471a}$.
